# A fatal plant toxicosis described more than a century ago is caused by a bacterial endophyte

**DOI:** 10.64898/2026.06.12.731819

**Authors:** Sandra Moreau, Manon Canal, Erin Hillairet, Jintao Ma, Max Crüsemann, Amélie Perez, Guillaume Marti, Marion Meyer, Aurélien Carlier

## Abstract

Gousiekte (‘quick disease’) is a fatal plant toxicosis affecting livestock in South Africa. Gousiekte is characterized by the sudden death of animals following the ingestion of leaves of several species of Rubiaceae, most notably *Vangueria pygmaea, Pavetta harborii, P. schumanniana* and *Fadogia homblei*. The toxin causing gousiekte is pavettamine, a hydroxylated polyamine pavettamine for which no biosynthetic pathway is known. Interestingly, all plants known to contain pavettamine also feature obligate endophytes of the *Burkholderiaceae* family, in particular belonging to the *Caballeronia* and *Paraburkholderia* genera. We identified a cluster of three genes conserved in all *Burkholderia s*.*l*. endophytic symbionts of plants containing pavettamine. Constitutive expression the *pavABC* gene cluster in a strain of *Paraburkholderia caledonica* isolated from leaves of *Fadogia homblei* resulted in detectable levels of pavettamine in cultures, and targeted gene deletions showed the involvement of all three genes in its biosynthesis. Genomes of important animal and human pathogens of the *Burkholderia pseudomallei* complex encode functional homologs of the *pavABC* genes, indicating a potentially unrecognized role of pavettamine in disease or complications from infections.

**Significance statement:** Gousiekte is a fatal disease of livestock caused by plants containing the toxin pavettamine, yet the origin of this compound has remained a mystery. We demonstrate that pavettamine is synthesized not by the plants themselves, but by their obligate bacterial endophytes via a conserved three-gene cluster, *pavABC*. Crucially, these genes were likely acquired via horizontal gene transfer from an ancestor of the *Burkholderia pseudomallei* complex, a group of significant human and animal pathogens. This discovery identifies the genetic basis of a potent toxin and suggests a previously unrecognized role for pavettamine in human infectious diseases. This work emphasizes the preeminent role of horizontal gene transfer and secondary metabolism for host-microbe interactions.

## Introduction

Gousiekte, literally “quick disease” in Afrikaans, is one of the most important plant toxicosis affecting livestock in southern Africa (1). Gousiekte is a significant and often fatal condition in ruminants, including cattle, sheep and goats, that occurs 4-8 weeks after the animals ingested the toxic plants (2). Some animals may exhibit outwards signs of congestive heart failure, but the main characteristic of the disease is sudden death from cardiac arrest sometimes triggered by physical exertion without prior signs of illness (3). Plants responsible for gousiekte poisoning include several Rubiaceae species such as *Fadogia homblei, Pavetta harborii, Pavetta schumanniana, Vangueria latifolia* (syn. *Pachystigma latifolium*), *Vangueria pygmaea* (syn. *Pachystigma pygmaeum)*, and *Vangueria thamnus* (syn. *Pachystigma thamnus*), although other species may also be involved (2, 4). These plants occur in the northeastern and central parts of South Africa and may be especially grazed when they sprout before the new grass appears, or in the late summer when grass becomes dry (5).

Gousiekte was formally described in 1923 (6), but this disease has proven difficult to diagnose because of varying or lack of early symptoms, differences in animal susceptibility, loss of toxicity as the plant dries, and the apparent seasonal toxicity of the plants (7–9). A chromatographic fraction from leaf extracts of *P. harborii* replicated the typical cardiomyopathy pathology of gousiekte when injected in goats, leading to the discovery of pavettamine (10). The structure of the toxin was solved in 2010, and shown to be an unusual hydroxylated polyamine (Figure 1) (9). Pavettamine purified from plant sources induced symptoms consistent with gousiekte in animal models and is assumed to be the sole causative toxin (8– 10). Pavettamine is suspected to inhibit protein synthesis because exposure of rat heart, kidney and liver cells to the toxin hinders the incorporation of radiolabeled phenylalanine into polypeptides. Consistent with this, cultures of rat H9c2 cells exposed to micromolar amounts of pavettamine exhibit mitochondrial damage, increased lysosomal activity and damage to the F-actin network (11).

**Figure 1.**
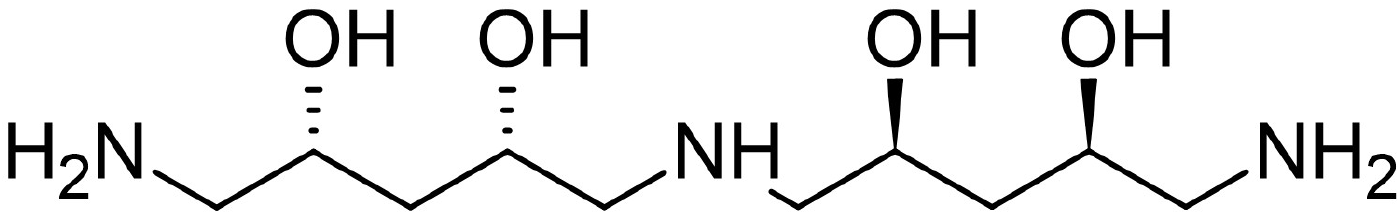
Structure of pavettamine. Adapted from Bode et al. 2010.

In addition to their association with gousiekte, *V. latifolia, V. macrocalyx, V. pygmaea*, and *V. thamnus* all present prominent colonies of bacterial endophytes in their leaves (7). These endophytic bacteria were later identified as belonging to the genus *Burkholderia* (4), later re-classified as *Paraburkholderia* and *Caballeronia* (12). The association between *Vangueria* and *Fadogia* species and *Burkholderia sensu lato* endophytes is highly specific and the bacteria appear to be vertically-transmitted (13). Moreover, leaf pavettamine content of *V. pygmaea* corelates with the abundance of endophytic bacteria, especially at the beginning of the growing season (14). Because of this systematic association, several authors have proposed a link between pavettamine and the presence of endophytes (4, 15, 16). Pavettamine has also been reported in several *Psychotria, Vangueria* and *Fadogia* species known to harbor obligate bacterial leaf endophytes that belong to the *Paraburkholderia* or *Caballeronia* genera (15). Although all specimen that tested positive for pavettamine also contained endophytic *Burkholderia s*.*l*., many more plant species associated with phylogenetically related bacteria did not contain detectable levels of pavettamine. There is currently no known biosynthetic pathway known for pavettamine, and whether the endophytes are responsible for the production of the toxic polyamine remains unclear.

Recent studies from our laboratories indicated that the genomes of vertically-transmitted, endophytic *Paraburkholderia* or *Caballeronia* species of Rubiaceae species often contain biosynthetic gene clusters (BGCs) responsible for the synthesis of various cytotoxic metabolites (e.g. aminocyclitols and non-ribosomal peptides) (17, 18). These biosynthetic gene clusters (BGCs) are often subject to horizontal gene transfer (HGT), and phylogenetically related species may harbor entirely different sets of BGCs. Conversely, unrelated species may harbor similar BGCs, indicating that functional complement does not always follow taxonomic boundaries (19, 20).

In the present study, we identified a 3-gene operon in several species of *Paraburkholderia* and *Caballeronia*, whose presence correlates with the content of pavettamine in host plants. Constitutive expression of the *pavABC* operon in a *Paraburkholderia* strain isolated from *Fadogia homblei* resulted in the production of pavettamine *in vitro*. Targeted deletion of each of the three genes resulted in loss of pavettamine production, demonstrating a direct role of *pavABC* operon in the production of the toxin and uncovering previously unknown metabolic intermediates. We also show that homologs of *pavABC* are widely distributed in leaf endophytic *Paraburkholderia* and *Caballeronia* species, as well as in strains of the *Burkholderia pseudomallei* complex, indicating potential unrecognized hazards linked to the production of pavettamine in these clinically important pathogens.

## RESULTS

### Identification of a gene cluster linked to pavettamine production

We reasoned that if endophytic *Caballeronia* or *Paraburkholderia* were responsible for the production of pavettamine, genes of the biosynthetic pathway would be conserved in the genomes of these endophytes, but otherwise rare in genomes of other *Burkholderiaceae* bacteria. Pavettamine has been detected in leaf extracts of *Psychotria kirkii, Pavetta schumanianna, Fadogia homblei* and *Vangueria pygmaea* (15). We have previously obtained metagenome-assembled genome (MAG) sequences of the *Caballeronia* and *Paraburkholderia* leaf symbionts of these species (17, 20, 21). We determined the core genome of a representative set of 337 *Burkholderiaceae* genomes, including 29 MAGs of leaf symbiotic endophytes. The core genome included sets of orthologous genes (orthogroups) with a representative in at least 80% of the genomes in the dataset. Next, we searched for orthogroups that did not belong to the core genome, but were present in 100% of the bacterial genomes assembled from samples of *P. kirkii, P. schumanianna, F. homblei* and *V. pygmaea*. We identified 48 orthogroups that fit these criteria (Table S4). We were unable to assign functional annotation for 25 of these orthogroups (COG category S or no hit). In addition, 8 groups contained proteins with putative functions without obvious link to enzymatic functions, such as transcriptional regulation (COG category K), translation and ribosomal structure (COG category J) and cytoskeleton (COG category U). Of the 15 remaining orthogroups, only one was annotated with functions related to amine metabolism and/or oxidation of substrates. The representative protein of OG0010878 (accession CCD35833) is annotated as a putative iron-sulfur cluster-binding protein, rieske family/carboxynorspermidine decarboxylase. The corresponding gene, (with locus tag in the *Ca*. B. kirkii UZHbot1 genome = BKIR_c114_0659) occurs in a putative operon with 2 other genes, with locus tags BKIR_c114_0658 (protein accession CCD35832) and BKIR_c114_0657 (protein accession CCD35831). The latter is also represented in the list of 48 accessory orthogroups that are conserved in all MAGs isolated from pavettamine-containing plants. This 3 gene putative operon is well conserved in all genomes or MAGs assembled from leaf samples of pavettamine-containing plants, although the genomic context surrounding the cluster is variable (Figure 2). In MAGs assembled from leaves of *V. pygmaea* and *F. homblei*, two of the most important plant species linked to gousiekte, the gene cluster is flanked on the 5’-end by a gene fragment showing homology to a truncated 2-epi-5-epi-valiolone synthase (EEVS). EEVS are key enzymes for the biosynthesis of C_7_-cyclitols such as streptol-glucoside and kirkamide, which are bioactive metabolites with herbicidal and insecticidal profiles isolated from other Rubiaceae plants involved in leaf symbiosis (21–23). Moreover, the cluster is flanked immediately upstream and downstream (within 500 bp) by IS elements of the IS5 family (figure 2). A homologous gene cluster is also present in the genome of *Ca*. Caballeronia hochstetterii, assembled from DNA isolated of leaves of *Pavetta hochstetteri*, a plant not yet analyzed for pavettamine content or known to be associated with gousiekte. However, the second gene of the operon presents an insertion causing a frameshift, likely inactivating the gene.

**Figure 2.**
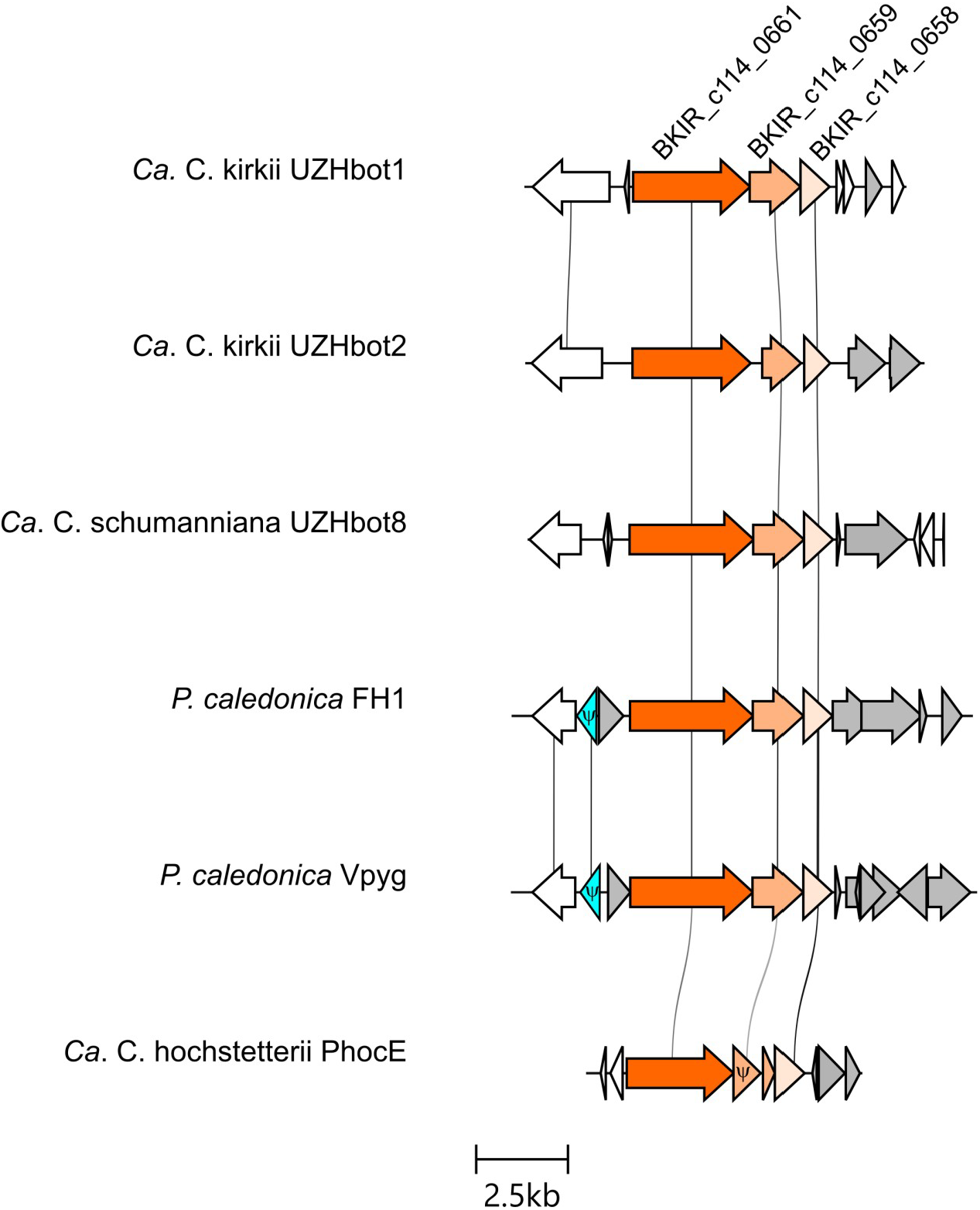
Gene structure of an operon conserved in the genomes of endophytes found in pavettamine-containing plants. Arrows colored in shades of orange denote bacterial candidate genes associated with pavettamine production in endophytes of *Psychotria kirkii* (*Ca*. C. kirkii UZHbot1 and UZHbot2), *Pavetta schumanniana* (*Ca*. C. schumanniana UZHbot8), *Fadogia homblei* (*P. caledonica* FH1), *Vangueria pygmaea* (*P. caledonica* Vpyg) and *Pavetta hochstetteri* (*Ca*. C. hochstetterii PhocE). Arrows colored in grey indicate IS sequences. Arrows in blue indicate gene fragments with homology to 2-epi-5-epi-valiolone synthases. Lines depict BLASTP pairwise identities > 50%. The Ψ symbols denote pseudogenes. The figure was created with clinker (62) and edited in Inkscape.

### Detection of pavettamine in cultures of *Paraburkholderia caledonica*

We isolated *Paraburkholderia caledonica* FH1 from leaves of *Fadogia homblei*, from which we were able to detect ions matching the expected m/z of derivatized pavettamine in leaf extracts by HPLC-MS/MS (Figure S1). We detected signals corresponding to the [M+H]^+^ and [M+Na]^+^ adducts of the polyamines putrescine and cadaverine, but not pavettamine in culture extracts of *P. caledonica* FH1 (Figure S1). However, biosynthetic genes may not be expressed under standard laboratory culture conditions (24– 26). To circumvent potential low expression of the cluster *in* vitro, we cloned the candidate *P. caledonica* SM1 ACR31T_39880-90 gene cluster on a broad-host range multicopy plasmid, downstream of a *P*_*EM7*_ synthetic promoter. Again, we were unable to detect ions corresponding to pavettamine in culture extracts of *P. caledonica* FH1, or *E. coli* transformed with the plasmid (raw data available on the Metabolights repository). To verify that the cloned genes were expressed in both bacterial hosts, we analyzed total cell extracts of *E. coli* pSEVA2313::ACR31T_39880-90 and *P. caledonica* FH1 pSEVA2313:: ACR31T_39880-90 by SDS-PAGE. Neither soluble nor insoluble protein fractions of *E. coli* displayed bands at the expected sizes for ACR31T_39890, ACR31T_39885 or ACR31T_39880. We detected distinct bands in the insoluble fractions of *P. caledonica* FH1 pSEVA2313::ACR31T_39880-90, but not in the soluble fractions (Figure S2). Proteins might form inclusion bodies upon overexpression in laboratory cultures, thereby potentially rendering the biosynthetic enzymes inactive. We repeated the experiment with *P. caledonica* FH1 pSEVA2313::ACR31T_39880-90, this time lowering the growth temperature to 16°C to prevent protein aggregation. We detected distinct peaks that corresponded to the [M+H]^+^ adducts of heptabenzoyl-pavettamine (Figure 3A). Furthermore, the MS^2^ spectra of the feature showed high similarity to the theoretical fragmentation pattern of heptabenzoyl-pavettamine (Figure S3).

**Figure 3.**
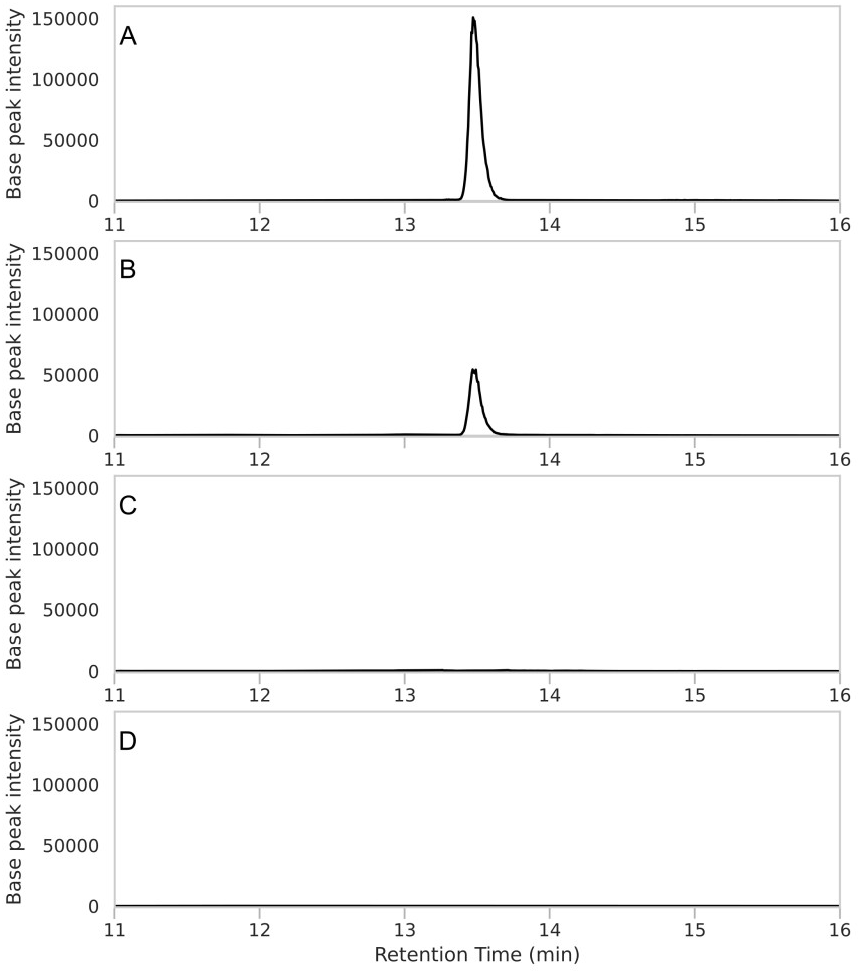
Detection of pavettamine in cultures of *P. caledonica* by HPLC-MS/MS. Leaf or bacterial culture extracts were derivatized with benzoyl-chloride as described in the main text. A. Extracted ion chromatogram with m/z = 980.3708 - 980.3802 of leaf extracts of *Fadogia homblei*., B. Cell extracts of cultures of *P. caledonica* FH1 pSEVA::*pavABC*; C. *P. caledonica* FH1 (wild-type), and D. Supernatant of cultures of *P. caledonica* FH1 pSEVA::*pavABC*.

However, we were unable to detect the metabolite in culture supernatants (Figure 3D), or in extracts of wild-type *P. caledonica* FH1 cultured in the same conditions (Figure 3C). These data demonstrate that pavettamine is produced by bacteria, and suggest that genes ACR31T_39890, ACR31T_39885 and ACR31T_39880 code for all or part of its biosynthetic pathway.

### Contribution of PavA, PavB and PavC to the biosynthesis of pavettamine

For the sake of simplicity, we will refer to the products of ACR31T_39890, ACR31T_39885 and ACR31T_39880 as PavA, PavB and PavC, respectively. PavA is a large (1119 AA) protein that is composed of 3 main domains. Alphafold and CATH analysis of the *Ca*. C. kirkii homolog reveals an N-terminal Rieske [2Fe-2S] iron-sulphur domain with sequence homology to naphthalene 1,2-dioxygenase α subunit, potentially involved in the incorporation of oxygen atoms into the backbone of the molecule. A central domain shows structural homology to an alanine racemase, and finally, a C-terminal domain is homologous to a domain found in lysine or ornithine decarboxylases (https://alphafold.ebi.ac.uk/entry/G4M3V6). PavA may therefore be involved in decarboxylation of an amino acid substrate such as lysine, as well as hydroxylation of the carbon backbone. PavB is a putative protein of unknown function containing a DUF3326 domain. Alphafold analysis indicates that the protein may contain a Rossman-like fold domain, found in proteins that bind cofactors like FAD, NAD+ or NADP+ (https://alphafold.ebi.ac.uk/entry/G4M3V5). Finally, PavC is a 264 AA protein containing a conserved JmjC-like cupin domain (https://alphafold.ebi.ac.uk/entry/G4M3V4). Members of this family are often involved in demethylation or hydroxylation reactions (27). To test whether *pavABC* is indeed directly involved in pavettamine biosynthesis, we constructed *P. caledonica* strain SM2, in which all 3 genes of the cluster have been deleted. We then introduced plasmids bearing each one of the following combinations: *pavABC, pavAB, pavAC* and *pavBC*. We cultured strain SM2 and derivatives under conditions permissive for pavettamine biosynthesis and analyzed bacterial extracts by HPLC-MS/MS after benzoyl-chloride derivatization. We could only detect pavettamine in extracts of *P. caledonica* SM2 expressing the entire *pavABC* operon (Figure 4P). The structure of pavettamine suggests that potential precursors donating 5-carbon chains with primary amines might possibly consists of lysine and/or cadaverine. We detected signals consistent with cadaverine in all extracts of *P. caledonica* SM2 and its derivatives (Figure 4A-E), as well as in extracts of *F. homblei* (Figure S1). Moreover, we detected signals for a possible biosynthetic intermediate N-(5-aminopentyl)pentane-1,5-diamine (i.e. compound 2, Figure 4) in extracts of bacteria expressing *pavABC* or *pavAB*, but not in strains lacking *pavA* or *pavB* (Figure 4F & H, Figure S4). Strains lacking *pavC* also accumulated a metabolite with a MS/MS signature resembling that of a possible dihydroxylated precursor of pavettamine (compound 3, Figure 4M, Figure S5). The same metabolite was also found in small quantities in extracts of bacteria expressing *pavABC* (Figure 4K). These data confirm that all three genes of the *pavABC* cluster participate to the biosynthesis of pavettamine.

**Figure 4.**
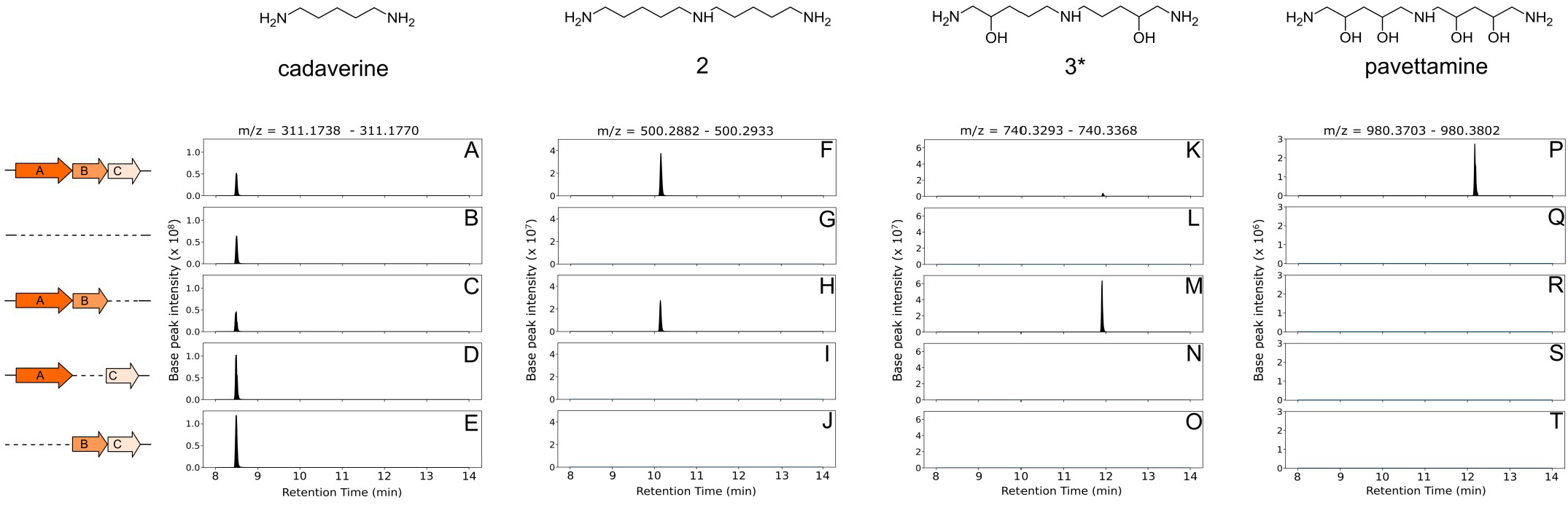
Detection of pavettamine and putative pathway intermediates in cultures of *P. caledonica* strains expressing all or a subset of the *pavABC* genes by HPLC-MS/MS. A to E: Extracted ion chromatograms corresponding to the expected m/z range of benzoyl derivatives of cadaverine as indicated in the top row. Chromatogram of culture extracts of: A, *P. caledonica* SM2 pSEVA2313::*pavABC*; B, *P. caledonica* SM2; C, *P. caledonica* SM2 pSEVA2313::*pavAB*; D, *P. caledonica* SM2 pSEVA2313::*pavAC*; E, *P. caledonica* SM2 pSEVA2313::*pavBC*. F to J: Extracted ion chromatograms corresponding to the expected m/z range of benzoyl derivatives of compound 2 as indicated in the top row. F. chromatogram of culture extracts of *P. caledonica* SM2 pSEVA2313::*pavABC*; G, *P. caledonica* SM2; H. *P. caledonica* SM2 pSEVA2313::*pavAB*; I. *P. caledonica* SM2 pSEVA2313::*pavAC*; J. *P. caledonica* SM2 pSEVA2313::*pavBC*. K to O: Extracted ion chromatograms corresponding to the expected m/z range of benzoyl derivatives of compound 3 as indicated in the top row. Note that the precise positions of the 2 hydroxyl groups are unknown. K, chromatogram of culture extracts of *P. caledonica* SM2 pSEVA2313::*pavABC*; L, *P. caledonica* SM2; M, *P. caledonica* SM2 pSEVA2313::*pavAB*; N, *P. caledonica* SM2 pSEVA2313::*pavAC*; O, *P. caledonica* SM2 pSEVA2313::*pavBC*. P to T: Extracted ion chromatograms corresponding to the expected m/z range of benzoyl derivatives of pavettamine as indicated in the top row. P, chromatogram of culture extracts of *P. caledonica* SM2 pSEVA2313::*pavABC*; Q, *P. caledonica* SM2; R, *P. caledonica* SM2 pSEVA2313::*pavAB*; S, *P. caledonica* SM2 pSEVA2313::*pavAC*; T, *P. caledonica* SM2 pSEVA2313::*pavBC*.

### Distribution of *pavABC* homologs in prokaryotes

To assess whether the *pavABC* cluster is present outside of the *Caballeronia* and *Parabukholderia* leaf symbionts of Rubiaceae, we searched for homologs in global protein databases. A preliminary search against the NCBI nr database revealed that homologs of all three genes of the cluster were present in 3668 publicly available genomes (Table S5). All of these hits occurred in genomes belonging to the *Burkholderia sensu lato*, i.e. species belonging to the *Burkholderia, Paraburkholderia* and *Caballeronia* genera. Specifically, a vast majority of the PavABC homologs were encoded in the genomes of the clinically-relevant *B. mallei* and *B. pseudomallei* species (3642/3668 genomes). Eight homologous clusters belonged to genomes of *B. humptydooensis*, which are environmental isolates of the *B. pseudomallei* species complex (28). Five clusters belonged to public genomes (MAGs) of leaf symbiotic *Caballeronia* species, including the symbionts of *Psychotria kirkii* and *Pavetta schumanniana*, while the taxonomy of the remaining 13 *Burkholderia* sp. hits was uncertain (Table S5). The *pavABC* cluster is present in a paraphyletic group of taxa, with at least 4 phylogenetically independent taxonomic groups (Figure 5). The *pavABC* cluster appears to be well represented in genomes of *B. pseudomallei*, with homologs in 1844/1879 (98.1%) of publicly available complete genomes. Similarly, 105/112 (93.7%) of genomes of *B. mallei* harbored *pavABC* homologs (Table S6). Phylogenetic analysis of homologs of PavB, PavC and the C-terminus of PavA show that despite a high degree of pairwise identity (>75% on average, Table S6), sequences of the *B. pseudomallei* complex and those of leaf symbiotic bacteria formed two distinct clades (Figure S6). Specifically, PavA homologs from leaf endophytic *Parabukholderia* and *Caballeronia* form a monophyletic clade, sister to that of the *B. pseudomallei* complex. However, *Paraburkholderia* and *Caballeronia* PavC sequences cluster basally to those of the *B. pseudomallei* complex (Figure S6C). This indicates a single origin of the *pavABC* operon in leaf symbiotic *Paraburkholderia* and *Caballeronia*, perhaps coopted from a direct ancestor of the *B. pseudomallei* complex.

**Figure 5.**
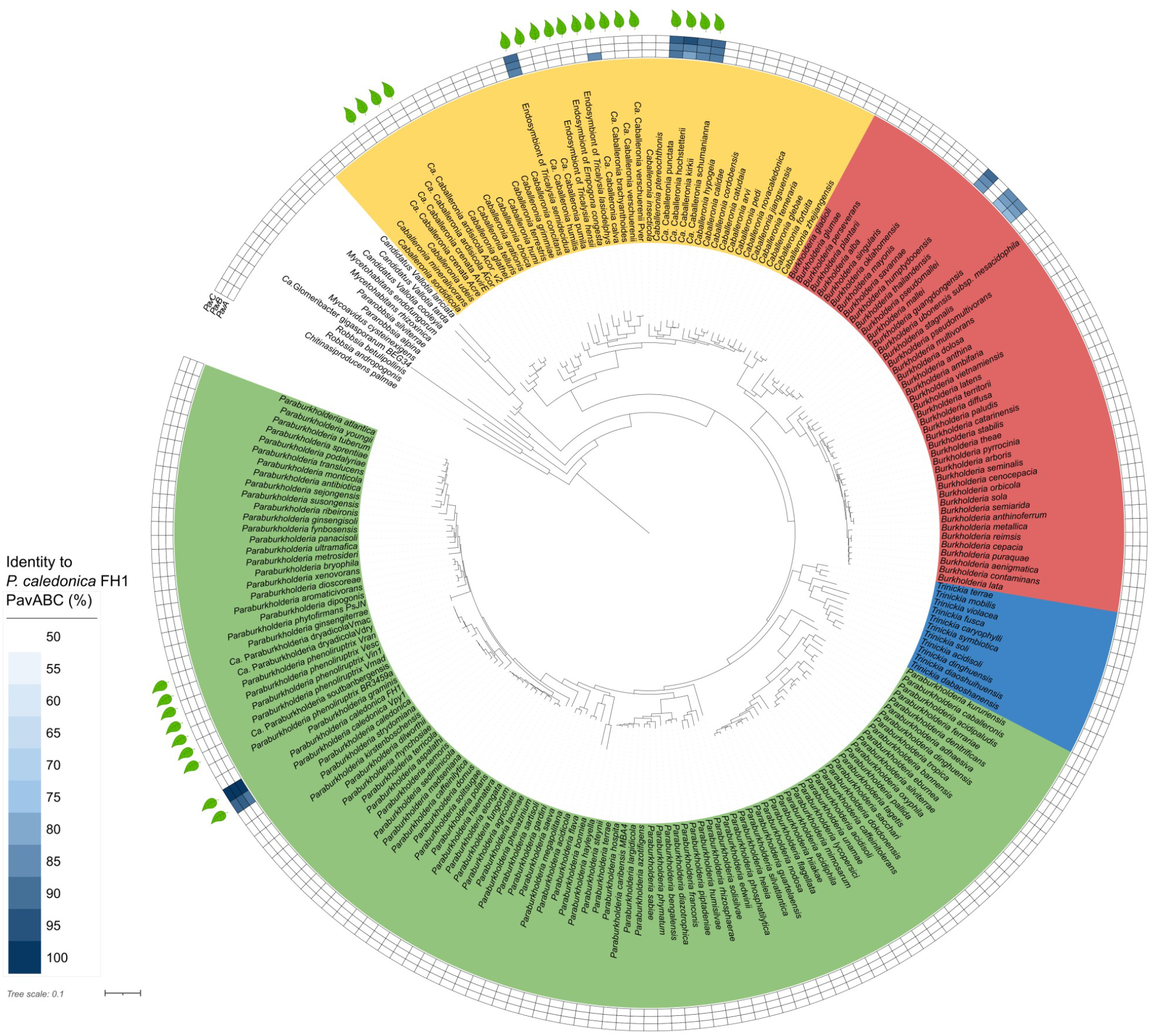
Distribution of *pavABC* homologs in representative genomes of *Burkholderia s*.*l*. The phylogenetic tree was created from a superalignment of core genes as detailed in the Materials and Methods. Colored ranges correspond to the following genera: *Burkholderia* (red), *Paraburkholderia* (green), *Trinickia* (blue) and *Caballeronia* (yellow). Outer rings denote the presence and % identity of PavABC homologs, as determined through an NCBI Blastp search with e-value < 10^−3^ and % identity > 50%. Leaf symbols denote genomes of leaf symbiotic bacteria of the Rubiaceae or Primulaceae.

### Functional analysis of the *pavABC* homologs of *Burkholderia pseudomallei*

Because the *pavABC* homologs of the *B. pseudomallei* complex formed a sister clade to those of leaf symbiotic *Burkholderia s. l*., we wondered whether homologs of *pavABC* in the *B. pseudomallei* complex encoded a functional pathway for pavettamine production. To test this, we introduced a synthetic gene cluster coding for the PavA, PavB and PavC homologs of *B. pseudomallei* K96243 into *P. caledonica* SM2 in an attempt to rescue pavettamine production in this strain. We were however unable to detect pavettamine or intermediates in culture extracts of that strain, perhaps due to weak expression of the synthetic construct (raw data available on the Metabolights repository). Expression of the synthetic *pavABC*_*Bpm*_ constructs in an *E. coli* protein expression platform resulted in the production of metabolites with identical features to the pavettamine intermediates 2 and 3, but not pavettamine itself (figure S7). This shows that all but the last steps of the pavettamine biosynthetic pathways may be reconstituted in *E. coli*, using PavABC protein sequences derived from *B. pseudomallei*.

## DISCUSSION

The link between gousiekte and leaf endophytes has intrigued scientists for over three decades (4, 7, 10). Beyond the obvious correlation between the presence of leaf nodules containing bacteria in several plants causing gousiekte, seasonal variation of pavettamine content and sensitivity to heat and dehydration are also reminiscent of the dynamics of peramine and loline accumulation in grasses colonized by endophytic fungi (29–31). Several lines of evidence indicate that the toxicity of pavettamine is an effective deterrent against herbivory. Perhaps the most compelling example is the report of a case in southern Zimbabwe. A herd of 27 African buffalos was captured at the Mutirikwe Recreational Park and transported to a new area in Mazowe, North of Harare. Three individuals died from apparent gousiekte symptoms shortly after being transported. Although herds of buffalos have lived in the area for several generations, it appears that the buffalos had been pushed into an unfamiliar area and likely grazed upon unfamiliar *Pavetta schumanniana*. Although *P. schumanniana* occurs widely in the area, disease had not been reported in wildlife before. Presumably, local herbivores learned to avoid grazing toxic plants (5). Together with the roles of several vertically-transmitted endophytic fungi in the production of allelopathic alkaloids (32) and our previous reports of bioactive peptides and carbasugars linked to leaf symbiosis, this indicates that defense against herbivory is a common function of hereditary plant symbioses.

Putative homologs of a cluster of 3 genes are present in all MAGs and genomes isolated from pavettamine-containing plants, but are otherwise rare in other *Paraburkholderia* and *Caballeronia* species. We named this cluster *pavABC* in the genome of *P. caledonica* SM1, likely forming a polycistronic operon. Because the species harboring homologs of *pavABC* form a polyphyletic group, it is likely that the gene clusters have been acquired multiple times through horizontal gene transfer. Consistent with this hypothesis, *pavABC* genes occur on plasmids in the two complete genomes of *Ca*. B. kirkii UZHbot1 and *P. caledonica* FH1 (pKIR01 and plasmid 2, respectively). Moreover, IS elements in the immediate vicinity of the operon are further indication of the possible role of HGT in the acquisition of the pavettamine biosynthesis pathway in leaf symbiotic bacteria. Homologous recombination between duplicated IS elements may mediate insertion and other structural variation in bacterial genomes (33, 34). Consistent with previous reports, cultures of *P. caledonica* isolates of *Fadogia homblei* do not produce pavettamine under standard laboratory conditions (35). This is likely because the biosynthetic genes are not expressed in these conditions, as expression of *pavABC* from a constitutive promoter is sufficient to induce production of the toxin. The *Ca*. C. kirkii orthologous proteins (locus tags BKIR_c114_0658, BKIR_c114_0659 and BKIR_c114_0661) figured among the 20 most abundant bacterial proteins in the *Psychotria kirkii* leaf nodule in a proteomics study, indicating that plant signals or cues may trigger expression of the pavettamine biosynthetic pathway (36).

Deleting any one of the three genes of the *pavABC* operon completely abolished the production of pavettamine in *P. caledonica*, demonstrating that PavA, PavB and PavC contribute directly to the biosynthetic pathway. However, because we were unable to detect pavettamine when *pavABC* were expressed in a heterologous system (*E. coli*), it is possible that other genes or co-factors participate to the biosynthesis. Because we detected compound 2 (N-(5-aminopentyl)pentane-1,5-diamine) in extracts of bacteria expressing both PavA and PavB, and that the C-terminal domain of PavA shows homology to a lysine decarboxylase, we propose that cadaverine and lysine might serve as the direct precursors of pavettamine. Indeed, we detected cadaverine in all samples of plant and bacterial cultures in which we also detected pavettamine (Figure 4). Deleting either *pavA* or *pavB* results in a complete loss of N-(5-aminopentyl)pentane-1,5-diamine (compound 2), suggesting that PavA and PavB both act simultaneously to decarboxylate lysine and condense cadaverine and lysine. However, we cannot rule out that other intermediates might be involved, that we fail to detect in our experiments. Alternatively, compound 2 may be a reaction byproduct instead of a true intermediate. Either PavA or PavB is also involved in the hydroxylation of intermediates. Expressing both PavA and PavB without PavC leads to the accumulation of a partially hydroxylated form of pavettamine, consisting of 2 instead of 4 hydroxyl groups. However, because we used mass spectrometry methods, we were unable to determine with certainty what positions of the carbon backbone are hydroxylated. It is likely given the symmetry of pavettamine that PavA/B hydroxylates carbons that are either in the *S* or *R* configuration, namely either positions 2 and 10, or 4 and 8. Finally, PavC likely catalyzes the hydroxylation at the remaining 2 carbon positions, yielding pavettamine. These results lead us to propose a hypothetical pathway for the synthesis of pavettamine (Figure 6). Understanding the precise contribution of each enzyme to pavettamine biosynthesis as well as the absolute configurations of the biosynthetic intermediates will be of high interest to develop new routes of stereospecific hydroxylation of polyamines and other aliphatic compounds.

**Figure 6.**
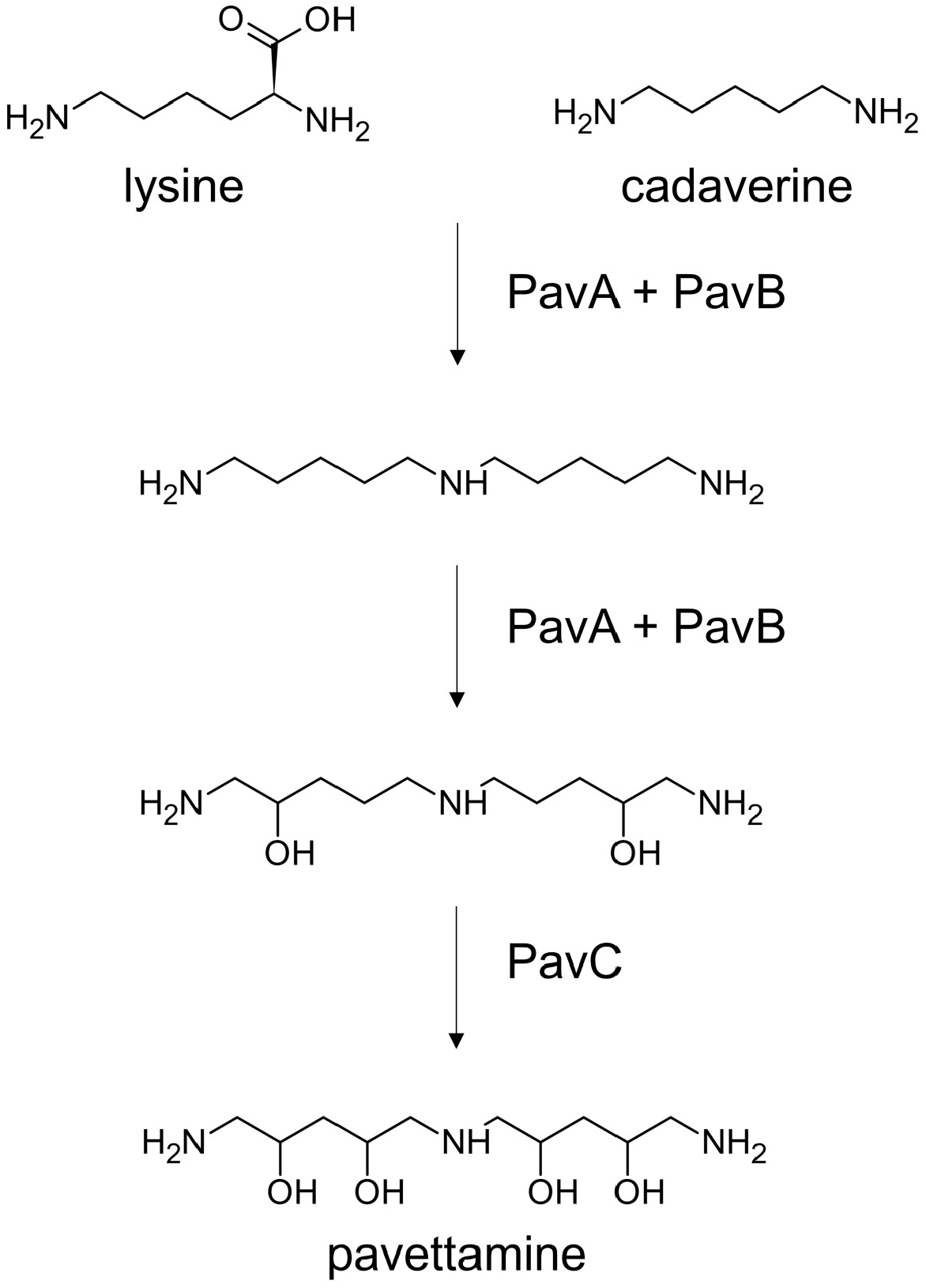
Proposed pathway of pavettamine biosynthesis. Likely precursors cadaverine and lysine are condensed and hydroxylated by the combined action of PavA and PavB, yielding intermediates 2 and 3. Note that the position of the hydroxyl groups of compound 3 is hypothetical. The final hydroxylation reaction is catalyzed by PavC, yielding pavettamine.

The finding of *pavABC* homologs in the *Burkholderia pseudomallei* group was unexpected. The *Burkholderia pseudomallei* group contains 5 species to date: *B. pseudomallei, B. mallei, B. thailandensis, B. humptydooensis* and *B. oklahomensis* (28). *B. thailandensis, B. humptydooensis* and *B. oklahomensis* are generally considered to be nonpathogenic soil-dwelling species, although some clinical cases of infection with these species have been reported (37, 38). *B. pseudomallei* and *B. mallei*, however, are known mostly as a cause of serious animal and human infections. *B. mallei* is the agent of glanders, a potentially fatal disease affecting horses, mules and donkeys, but rarely humans (39). *B. mallei* is often considered to be essentially a clone of *B. pseudomallei* that has become adapted to equines hosts (40).

*B. pseudomallei* is a major human pathogen, causing the life-threatening disease melioidosis (41). Melioidosis results from environmental exposure to *B. pseudomallei* by inhalation, inoculation or ingestion (42, 43). Mortality following infection is high, from 10% to 40%, and is compounded by limited treatment options due to broad antimicrobial resistance (44). As a result, *B. pseudomallei* is classified as a Risk Group 3 agent in several countries where it is not endemic, e.g. in countries of the European Union, the UK, Canada, and as a Tier 1 select agent by the US Center for disease control. *B. pseudomallei* is usually found in wet, acidic soil, as well as in the rhizosphere of grasses and fecal matter of grazing animals in South East Asia and Australia (45). *B. pseudomallei* employs a broad array of virulence factors to cause infection, gain intracellular entry and evade the host immune system. For example, the genome of *B. pseudomallei* encodes three type III (T3SS) and six type VI (T6SS) secretion systems that deliver protein effectors to host cells, promote intracellular survival and competition with resident microbiota (41). In addition to these, *B. pseudomallei* secretes the *Burkholderia* lethal factor 1 (BLF-1, a protein shown to modify the human translation initial factor elF4A) (46), as well as several cytotoxic specialized metabolites (e.g. malleilactone and malleicyprols) (47, 48). Expression of *B. pseudomallei* PavABC proteins in *E. coli* successfully reconstituted the first and second steps of the hypothetical pathway of pavettamine synthesis, indicating that *B. pseudomallei* PavA and PavB enzymes are fully functional. The fact that we were unable to detect pavettamine in that setting suggests that our synthetic *B. pseudomallei* PavC construct is either non-functional, may require co-factors that are absent in *E. coli*, or is not expressed or folded correctly in this system. Regardless, our data provide strong evidence that the *pavABC*_*Bpm*_ operon of the *B. pseudomallei* complex encodes a *bona fide* pavettamine biosynthetic pathway.

The role of pavettamine in the ecology of the *B. pseudomallei* complex is unknown. Pavettamine could perhaps act in concert with the *Burkholderia* lethal factor 1 (BLF-1) to inhibit translation in hosts (46). Expression of *B. pseudomallei* K96243 *pavA* (BPSS1453), pavB (BPSS1454) and *pavC* (BPSS1455) has been detected in 43/82 conditions tested in a large microarray experiment, although notably not in mouse macrophage or mouse lung infection (49). Furthermore, BPSS1453 and BPSS1455 are downregulated by an average of 19-fold when cultured in human plasma versus conditions mimicking soil (50). This indicates that pavettamine may play a role during the free-living stages of *B. pseudomallei* lifecycle, or perhaps during the early phases of infection. Additionally, whether pavettamine remains intracellular or is secreted remains unanswered.

In conclusion, our study sheds light on what is probably the clearest example yet of a plant toxicosis mediated by a bacterial endophyte. Indeed, we demonstrate that bacteria of the *Paraburkholderia* and *Caballeronia* genera are solely responsible for the synthesis of pavettamine, the toxin causing gousiekte. Furthermore, studies into the molecular mechanisms leading to gousiekte symptoms and death have so far been hindered by the difficulty in securing large quantities of the toxin. Extraction and purification of pavettamine from plants requires large quantities of leaves, as the overall yield is quite poor. For example, one study reports yields of 2.7 mg of purified pavettamine per kg of dried plant material (11). In addition, pavettamine may be synthesized chemically, but requires over 25 steps to achieve enantiomeric purity. Our study thus offers new possibilities for the production of pavettamine from bacterial cultures, and new studies towards better understating its mode of action.

## Materials and methods

### Strains and culture conditions

Strains and plasmids used in this study are reported in Table S1. *P. caledonica* FH1 was isolated from leaves of *Fadogia homblei* collected in the Gauteng province (South Africa) in Spring 2020 on tryptic soy agar (TSA). The strain was collected in compliance with local and international regulations, and was shared between laboratories involved in the study under South African export permit n° BABS-1139 (due diligence declaration available at https://absch.cbd.int/en/database/CPC/ABSCH-CPC-FR-294070-1). All *E. coli* and *P. caledonica* strains were grown in LB or tryptic soy agar (TSA, Oxoid) medium at 37°C and 28°C, respectively with or without kanamycin 50µg/mL. Liquid cultures were done in LB or TSB, kanamycin 50µg/mL, 28°C. All chemicals were purchased from Merck unless otherwise indicated. *P. caledonica* SM1 is a spontaneous nalidixic acid-resistant strain obtained after serial passaging of a clone of *P. caledonica* FH1 on TSA medium containing increasing concentrations of nalidixic acid, from 10 µg/mL to 30 µg/mL.

### Genome sequencing and annotation

Genomic DNA from overnight cultures of strain SM1 was extracted using CTAB and phenol-chloroform followed by isopropanol precipitation (51). Contaminating RNA was digested using RNAse A (Qiagen) at 2 mg/mL, 5 µL per 100 µL extract and incubated for 1h at 37°C. DNA was further purified using Sera-Mag Select magnetic beads (Cytiva) according to manufacturer recommendations. DNA quality was checked using 1% agarose gel electrophoresis and quantification was done using a Qubit 3 instrument and kit (Thermo Fisher Scientific). DNA was stored at -20°C prior to further analysis. Sequencing libraries were prepared using the Oxford Nanopore Technologies Native Barcoding kit SQK NBD 112.4 and sequenced on an Oxford Nanopore P2solo instrument using a R10.4 flow cell. Base calling was done with Dorado v.0.9 in GPU mode on a Linux PC equipped with an NVIDIA A6000 graphics card and 48 Gb of on-board memory. Reads were filtered with chopper (52) with default settings with the following settings: -q 10 -l 1500 --headcrop 10 --tailcrop 10. Filtered reads were assembled on a Linux server running Debian 5.16.11-1 (64 cores and 504 Gb of RAM) with Canu v2.2 (53) with the following options : useGrid=False, corOutCoverage=100, genomeSize=6.3m, enableOEA=false. Contigs were circularized using an ad-hoc python script. The genome was annotated with the NCBI Prokaryotic Genome Annotation Pipeline v. 6.3 (54) and deposited in NCBI Genbank under accessions CP195752-CP195755. IS sequences were predicted with ISEscan with default settings (55).

### Comparative genomics

The proteome sequences of a 308 representative set of *Burkholderiaceae* species were downloaded using the NCBI datasets tool, with the “taxon 119060 --annotated --representative --assembly-source ‘RefSeq’ --include protein” (56). The dataset was complemented with the predicted proteomes of metagenome-assembled genome (MAG) sequences of leaf symbiotic *Paraburkholderia* and *Caballeronia* species obtained from (21). A list of accession numbers is available in Table S3. Predicted pseudogenes were excluded from the analysis. Predicted proteomes were compared using OrthoFinder v2.5.2 (57) with default settings. Core orthogroups were defined as containing genes from >80% of the genomes in the analysis. The accessory genome of a given MAG or strain was defined as the set of orthogroups with at least two sequences, and not belonging to the core genome. Candidate genes involved in pavettamine biosynthesis were selected based on accessory orthogroups with members in all genomes of endophytes of certified pavettamine-containing plant species, i.e. *Vangueria pygmaeae, Fadogia homblei, Pavetta schumanniana* and *Psychotria kirkii*. One representative sequence per candidate orthogroup was selected from the *Ca*. B. kirkii UZHbot1 publicly available assembly, and used as a query for a search against the EggNOG database to provide functional annotation (58). Global BLASTP searches against the NCBI nr database were conducted with cblaster (59), with --max_evalue = 0.001 and --min_identity = 50. To generate a phylogenetic tree of the *Burkholderiaceae*, AA sequences of core genes identified by Orthofinder were aligned using MAFFT in --auto mode. Alignments were trimmed and concatenated and used to create a super tree using FastTree with the -JTT model (60). Newick files were visualized and annotated in iTol (61). Proteomes of the *Burkholderia pseudomallei* group were downloaded with NCBI datasets v.18.0.2 with the following parameters “taxon 111527 --annotated --assembly-source ‘RefSeq’ -- include protein”. The presence of homologs of the PavABC proteins was assessed for each proteome using NCBI blastp v2.11.0+ with the sequences of PavABC proteins of *P. caledonica* SM1 as queries. Only hits with e-value < 1e-6 were reported. Visualization of gene cluster alignments was done with clinker (62).

### Bacterial genetics and cloning

A 5973 bp DNA fragment containing the *P. caledonica* SM1 genes ACR31T_39880-90 was obtained by PCR using primers Pcal_pavettamine_R-XbaI and Pcal_pavettamine_F-BamHI (Table S2). The 5.9 kb PCR product was digested with BamHI and XbaI (New England Biolabs), purified using agarose gel electrophoresis and ligated into plasmid pSEVA2313 to yield pSEVA2313::*pavABC*. Ligation products were transformed into *E. coli* Top10 using a standard heat-shock protocol. Transformants were selected on LB plates containing kanamycin 50 µg/mL. One transformant was selected for following experiments and the plasmid construct was validated by sequencing in-house using ONT Nanopore v14 Rapid barcoding kit and R10.4.1 flow cells (63).

Plasmid pSEVA2313::*pavABC* was introduced into *P. caledonica* strains by electroporation. Deletion mutants were produced in a *P. caledonica* SM1 background by homologous recombination using a previously published protocol (64). Briefly, ∼500 bp fragments flanking the operon were amplified by PCR, using primers designed to feature a BsaI restriction sites (Table S2). DNA of plasmid pSNW2 was amplified by PCR and ligated with the insert by Golden Gate cloning. Ligation products of the inserts and vector were introduced into electrocompetent *E. coli*. The construct was validated by Nanopore ONT sequencing as above. Plasmids were introduced into *P. caledonica* SM1 by conjugation. Conditionally-replicating plasmid pQURE6 harboring a I-SceI nuclease gene was introduced by electroporation into kanamycin-resistant merodiploid clones. Colonies with gene replacement events were screened for loss of kanamycin resistance. Clone *P. caledonica* SM2 was selected for downstream analysis, and its genome sequence was obtained by Oxford Nanopore sequencing using R10.4 chemistry on an ONT P2 solo instrument, confirming deletion of the targeted locus and an absence of spurious mutations. For *in trans* partial complementation experiments, plasmids harboring *pavABC* genes were amplified by PCR from plasmid pSEVA2313::*pavABC* using PrimeSTAR Max DNA Polymerase (Takara Bio) with primers listed in Table S2. Resulting PCR products were digested with DpnI to remove template DNA and circularized with T4 ligase (New England Biolabs) overnight at 16°C. Ligation products were transformed into *E. coli* by electroporation, and clones selected on LB medium supplemented with kanamycin (50 mg/L). Plasmid DNA was isolated from selected clones and sequenced by ONT Nanopore sequencing as above. Plasmids were then introduced into *P. caledonica* SM2 by electroporation. Transformants were selected on TSA medium supplemented with nalidixic acid 30 µg/mL and kanamycin 50 µg/mL. A synthetic operon coding for the PavA, PavB and PavC homologs of *B. pseudomallei* K96243 was designed using the genome sequence (NCBI accession ASM95928v1) as a template. Each CDS sequence was optimized to remove tandem repeats and avoid high %GC regions using the GeneArt optimization tool. The resulting 5.6 kb fragment was synthesized by Life Technologies SAS (Courtaboeuf, France) and cloned into the pSEVA2313 broad host range plasmid vector or into plasmid pLS286. Whole plasmid constructs were sequenced with ONT Nanopore as above and deposited in NCBI Genbank (accessions PZ017930 and PZ017931). Plasmid pSEVA2313::*pav*_*Bpm*_*ABC* was introduced into *P. caledonica* SM2 by electroporation as above. Plasmid pLS386::*pav*_*Bpm*_*ABC* was introduced into *E. coli* ER2566 by chemical transformation using standard protocols.

### Optimization of protein production

Unless otherwise mentioned, *E. coli* strains were cultured in 10 ml LB containing kanamycin 50µg/ml at 37°C for 24h or at 28°C for 48h. *P. caledonica* strains were cultured in TSB medium at 28°C for 48h. Cells of *E. coli* pSEVA2313::*pavABC* or *P. caledonica* pSEVA2313::*pavABC* were harvested by centrifugation at 10000 x g for 5 min and lysed using BugBuster protein extraction reagent (Millipore) and lysonase (Millipore) according to the suppliers’ recommendations. The mixture was then centrifuged for 20 min at 20000 x g at 4°C and the supernatant separated from the pellet. Proteins present either in the supernatant (soluble proteins) or in the pellet (insoluble proteins) were denatured in 2x Laemmli sample buffer for 10 min at 95°C. Samples were loaded on precast SDS-PAGE gels (4-15%; Biorad, Hercules, CA, USA) and stained with InstantBlue (Abcam). Cultures of *E. coli* ER2566 harboring pLS386 and derivatives were grown in 100 mL of Terrific Broth (TB) medium supplemented with ferric citrate (100 mg/L), ferrous sulfate heptahydrate (100 mg/L), ferric ammonium citrate (100 mg/L) and L-cysteine (1 mM), and 0.1 mM of isopropylthio-β-galactoside (IPTG).

### HPLC-MS analysis of pavettamine

Leaves of *Fadogia homblei* were previously collected at Roodeplaat, near Pretoria, South Africa (21) (voucher PRU 128010). After collection, fresh leaves were stored in airtight 50 mL polypropylene tube and immediately frozen at -80°C. Approximately 150 mg of frozen leaf material were ground in liquid nitrogen with mortar and pestle. Derivatization with benzoyl chloride was adapted from (35). Briefly, 2 mL of an aqueous solution of perchloric acid (5% v/v) were added to the sample, mixed thoroughly and stored on ice for 30 min. Debris were removed by centrifugation at 20 000 x g for 20 min. Approximately 250 µL of supernatant were added to a 2 mL borosilicate tube containing 1.5 mL of 2N NaOH and 2 µL of a diaminoheptane (100 mM) standard. After several minutes at room temperature to equilibrate, 20 µL of reagent-grade benzoyl chloride were added, mixed by vortexing and left at room temperature for 20 min. Derivatized amines were extracted with 4 mL of HPLC-grade diethyl ether. The organic phase was washed twice with an equal volume of ultrapure water, and evaporated to dryness under a stream of nitrogen. Samples were kept at -20°C until analysis.

Cultures of *P. caledonica* strains were grown in 500 mL flasks containing 100 mL of TSB medium supplemented with kanamycin 50 µg/mL as appropriate. Cultures were incubated at 16°C with shaking (120 rpm) for 96h. Cultures were centrifuged at 10000 x g for 10 min to separate cells and supernatant. Cell pellets were washed once with 4.5 mL of NaCl 0.4% (w/v), resuspended in 5 mL of 2N NaOH and stored on ice. Cells were lysed on ice by sonication with 6 × 30s bursts. Per 2 mL of lysate, 20 µL of diaminoheptane standard (100 mM) were added followed by 20 µL of benzoyl chloride. Reactions were left at room temperature for 20 min, and derivatized amines were extracted with 4 mL of diethyl ether as above. Samples were dried and extracts were accurately weighed prior to diluting in solvent. Ultra-high-performance liquid chromatography-high-resolution MS (UHPLC-HRMS) analyses were performed on a Q Exactive Plus quadrupole (Orbitrap) mass spectrometer, equipped with a heated electrospray probe (HESI II) coupled to a U-HPLC Ultimate 3000 RSLC system (Thermo Fisher Scientific, Hemel Hempstead, UK). Separation was done on a Luna Omega Polar C18 column (150 mm × 2.1 mm i.d., 1.6 μm, Phenomenex, Sartrouville, France) equipped with a guard column. The mobile phase A consisted of water with 0.05% formic acid (FA), and the mobile phase B was acetonitrile with 0.05% FA. The solvent gradient was 0 min, 100% A; 1 min, 100% A; 22 min, 100% B; 25 min, 100% B; 25.5 min, 100% A; and 28 min, 100% A for re-equilibration. The flow rate was 0.3 mL/min, the column temperature was set to 40°C, the autosampler temperature was set to 5°C, and the injection volume was fixed to 5 μl. Mass detection was performed in positive ionization (PI) mode at resolution 35,000 power full width at half-maximum (fwhm) at 400 m/z] for MS1 and 17,500 for MS2 with an automatic gain control (AGC) target of 1 × 10^6^ for full scan MS1 and 1 × 10^5^ for MS2. Ionization spray voltages were set to 3.5 kV, and the capillary temperature was kept at 256°C. The mass scanning range was m/z 100-1500. Each full MS scan was followed by data-dependent acquisition of MS/MS spectra for the six most intense ions using stepped normalized collision energy of 20, 40, and 60 eV. Raw data were processed with MZmine version 4.6.1 for mass signal extraction between 100 and 1,500 Da in positive mode (65). Noise level was set to 1.0E3. The m/z tolerance was set to 5 ppm. For chromatogram building, minimum absolute height was set to 1.0E4 and minimum intensity for consecutive scans = 1.0E3. Local minimum feature resolver was used with minimum absolute height = 1.0E5, peak duration = 0.00 - 10.00, minimum scans = 3. Theoretical m/z of M^+^, [M+H]^+^, and [M+Na]^+^ adducts of benzoylated pavettamine and intermediates were predicted in ChemSketch v.2024.1.4 (ACD/Labs). Prediction of MS/MS fragmentation patterns was done by searching SMILES structures using the CFM-ID Web Server Spectra prediction utility (66). All experimental HPLC MS/MS data were deposited in the Metabolights database (www.ebi.ac.uk/metabolights) under accession MTBLS13921. Reviewer link : https://www.ebi.ac.uk/metabolights/reviewer3579ece2-df57-49c1-b021-dd414789c9cf

## ACKNOWLEDGMENTS

We wish to thank Laurent Sauviac (LIPME) for sharing *E. coli* strains and plasmids used in this study, as well as the SEVA team for making useful plasmid resources available. We also wish to thank Karine Laroucau (ANSES, Maisons-Lafort, France) for helpful comments on the manuscript.

## DATA AVAILABILITY

The genome sequence of *Paraburkholderia caledonica* SM1 is available from NCBI Genbank with accession numbers CP195752-CP195755. Experimental HPLC-MS/MS data and associated metadata are available at https://www.ebi.ac.uk/metabolights/MTBLS13921.

Sequences of synthetic constructs used in this study are available in NCBI Genbank with accessions PZ017930 and PZ017931.

