## supplementary figures for "A fatal plant toxicosis described more than a century ago is caused by a bacterial endophyte"

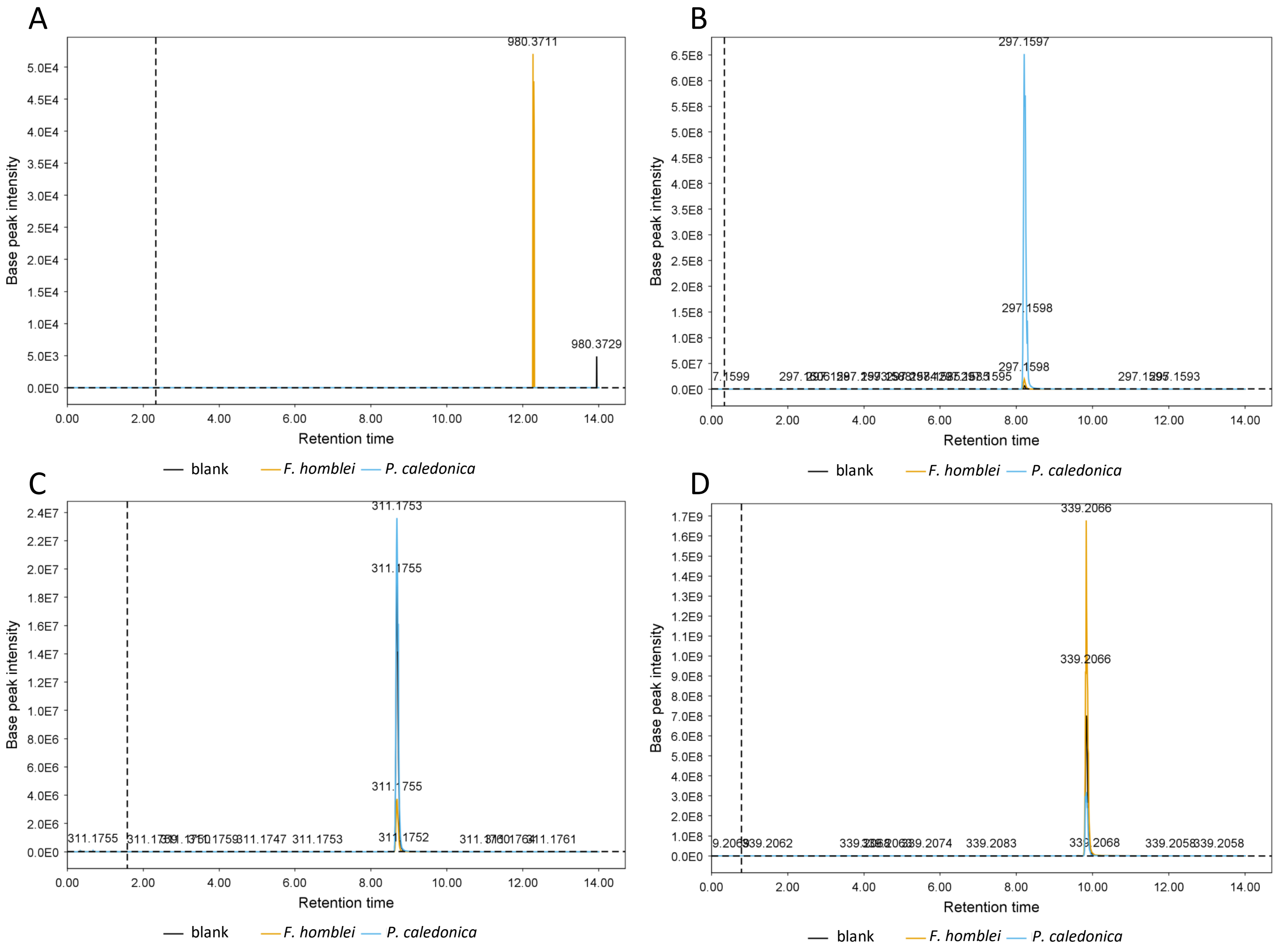


**Figure S1. Detection of polyamines by HPLC-MS/MS.** Amines were derivatized with benzoyl chloride, extracted with diethyl ether and separated by reverse-phase HPLC MS/MS with data acquisition in positive mode. Extracted ion chromatograms were produced using Mzmine 4.6.1, using the predicted mass-to-charge (m/z) ratios of the M+, [M+H]+ and ]M+Na]+ adducts (with tolerance set at +/- 5 ppm). Only chromatograms corresponding to the [M+H]+ adducts are shown. A, Extracted ion chromatogram with set m/z = 980.3752 ± 5 ppm (heptabenzoyl pavettamine). B, Extracted ion chromatogram with set m/z = 297.1597 ± 5 ppm (benzoyl putrescine).  C. Extracted ion chromatogram with set m/z = 311.1754 ± 5 ppm (benzoyl cadaverine). D. Extracted ion chromatogram with set m/z = 339.2067 ± 5 ppm (benzoyl diaminoheptane, internal standard). Yellow lines: *Fadogia homblei* leaf extracts; Blue lines: *Paraburkholderia caledonica* culture extracts; Grey lines: extraction blank.

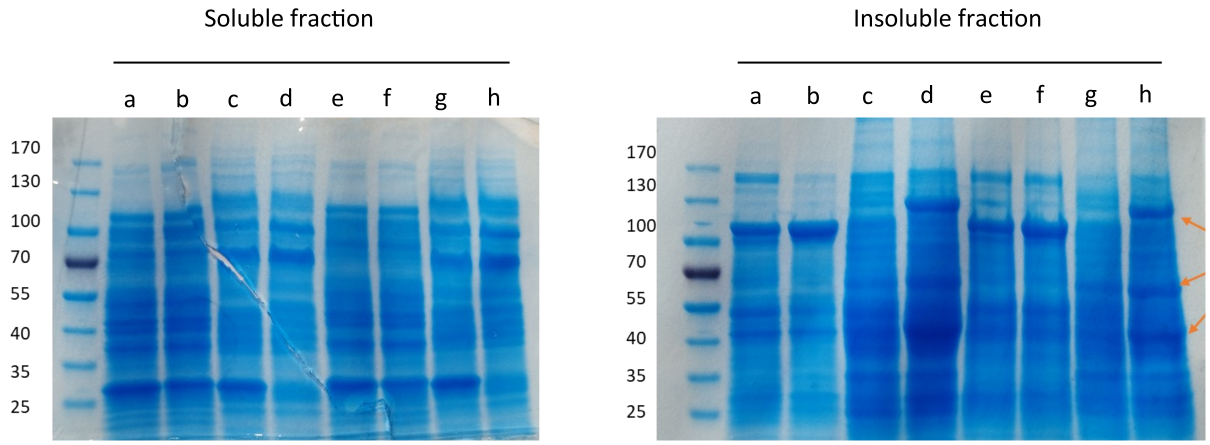


**Figure S2. SDS-Page analysis of cell extracts of *E. coli* Top10 and *P. caledonica* FH1 harboring pSEVA2312::*pavABC* and pSEVA2313::*gfp*.** Cultures were grown at 21°C with shaking for 48h. Bacteria were harvested by centrifugation and lysed with BugBuster reagent. Extracts were centrifuged to separate the soluble and insoluble fractions. *a, E. coli* pSEVA2313*::gfp* grown in LB; b, E. coli pSEVA2313::pavABC in LB; c. *P. caledonica* FH1 pSEVA2313::*gfp* in LB; d. *P. caledonica* FH1 pSEVA2313::*pavABC* in LB; *e, E. coli* pSEVA2313*::gfp* grown in LB; f, E. coli pSEVA2313::pavABC in TSB; g. *P. caledonica* FH1 pSEVA2313::*gfp* in TSB; h. *P. caledonica* FH1 pSEVA2313::*pavABC* in TSB. Orange arrows indicate the expected size of PavA, PavB and PavC.

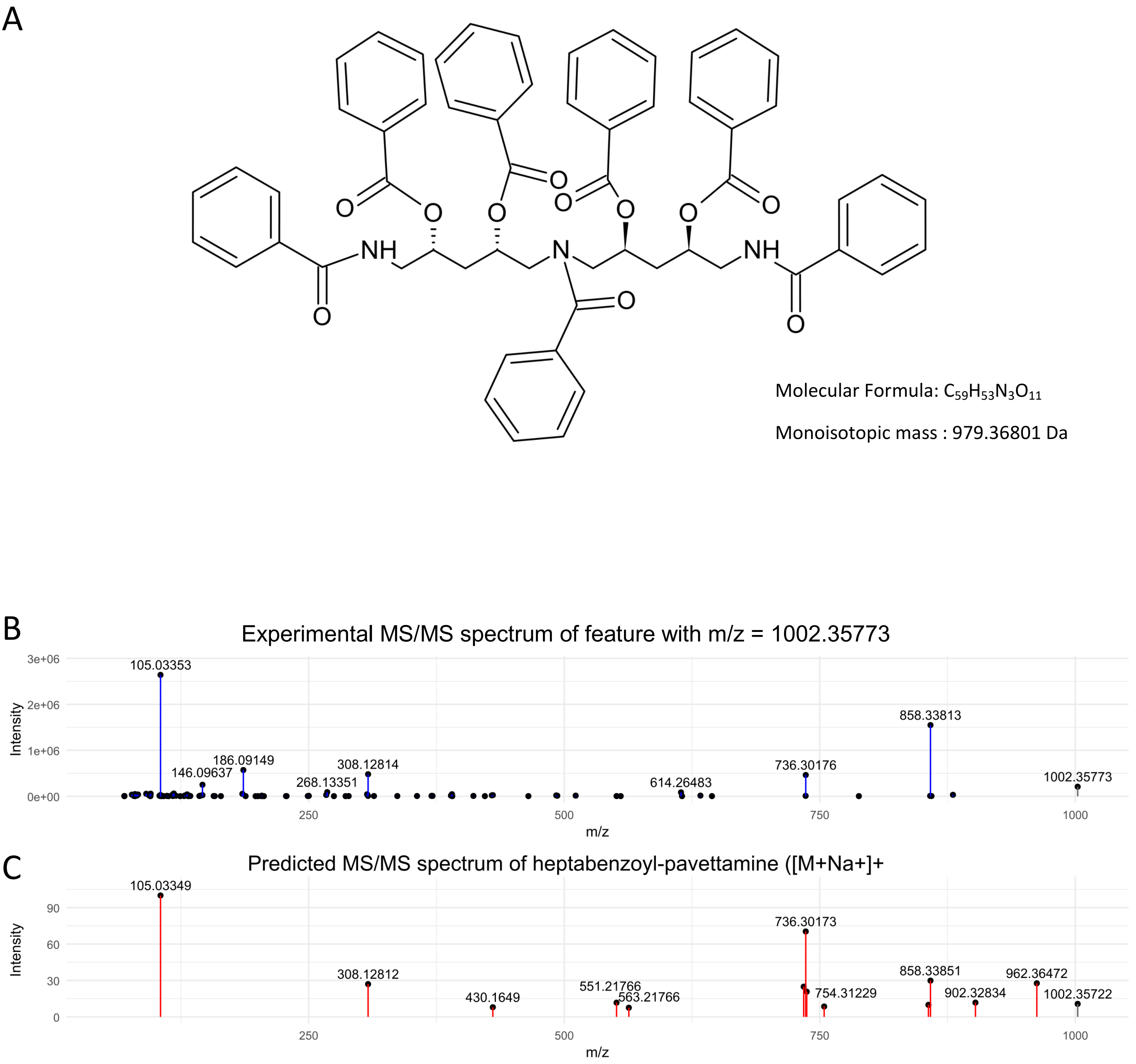


**Figure S3. MS/MS spectra of benzoylated features detected in cell extracts of *P. caledonica***. A. Predicted structure and molecular properties of heptabenzoyl-pavettamine. B. MS/MS profile of a feature with m/z= 1002.3569 detected in derivatized extracts of cultures of *Paraburkholderia caledonica* FH1 pSEVA2313::*pavABC* grown at 16°C in LB medium. C. Predicted MS/MS spectrum of heptabenzoyl-pavettamine determined using the CFM-ID web server with the SMILES notation: O=C(OC(CN(CC(OC(=O)c1ccccc1)CC(OC(=O)c1ccccc1)CNC(=O)c1ccccc1)C(=O)c1ccccc1)CC(OC(=O)c1ccccc1)CNC(=O)c1ccccc1)c1ccccc1). Peaks in grey indicate precursor ions. Experimental MS data were analyzed in MZmine 4.6.1. Spectra data were exported and processed with the ggplot2 R package to generate the plots.

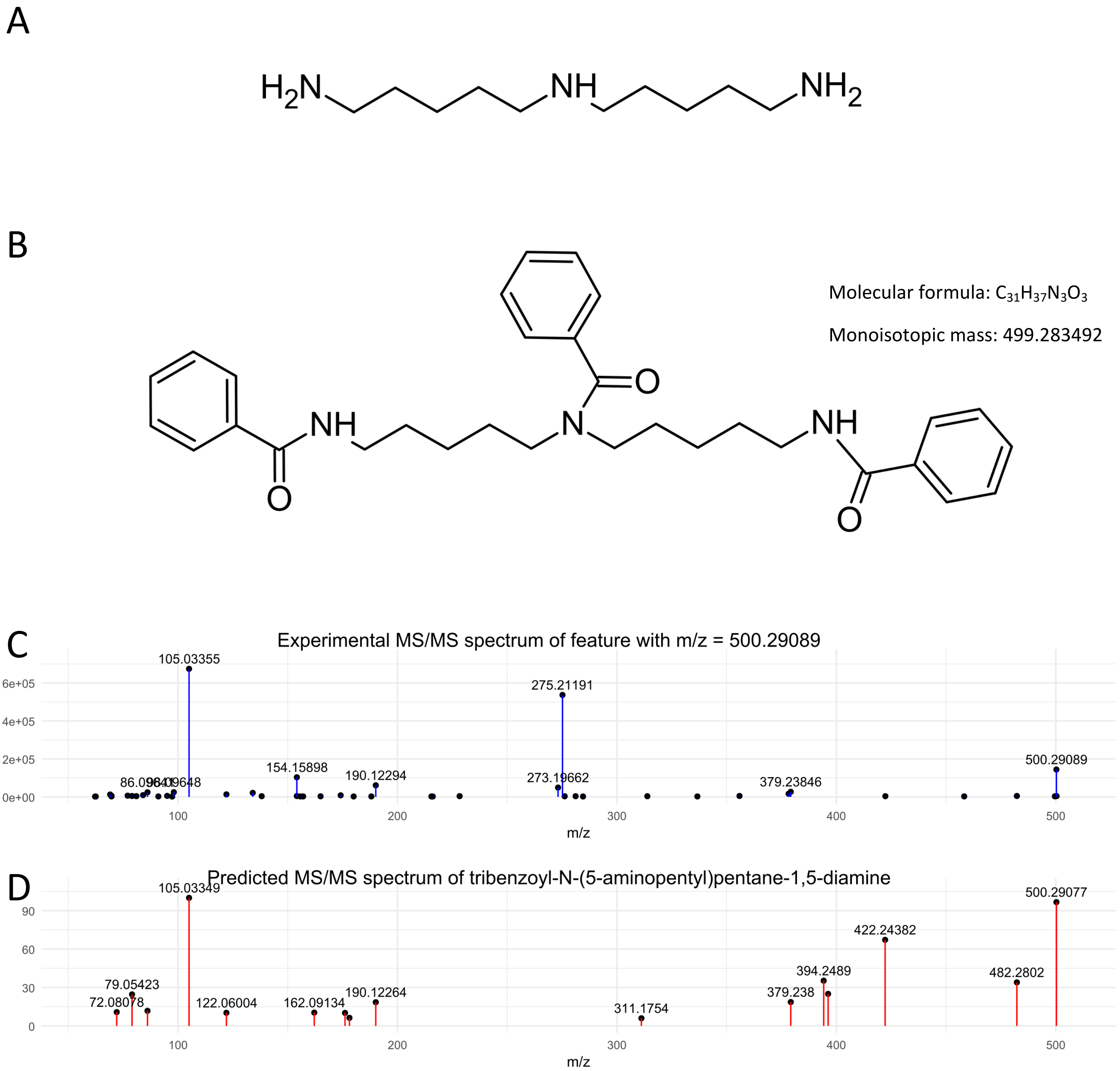


**Figure S4. MS/MS spectra of benzoylated features detected in cell extracts of *P. caledonica***. A. Predicted structure and molecular properties of tribenzoyl-bis(5-aminopentyl)amine. B. MS/MS profile of a feature with m/z= 500.2909 detected in derivatized extracts of cultures of *Paraburkholderia caledonica* SM2 pSEVA::*pavAB* grown at 16°C in LB medium. C. Predicted MS/MS spectrum of tribenzoyl-N-(5-aminopentyl)pentane-1,5-diamine determined using the CFM-ID web server with the SMILES notation: O=C(N(CCCCCNC(=O)c1ccccc1)CCCCCNC(=O)c1ccccc1)c1ccccc1 and collision energy set at 20 eV. Experimental MS data were analyzed in MZmine 4.6.1. Spectra data were exported and processed with the ggplot2 R package to generate the plots.

A

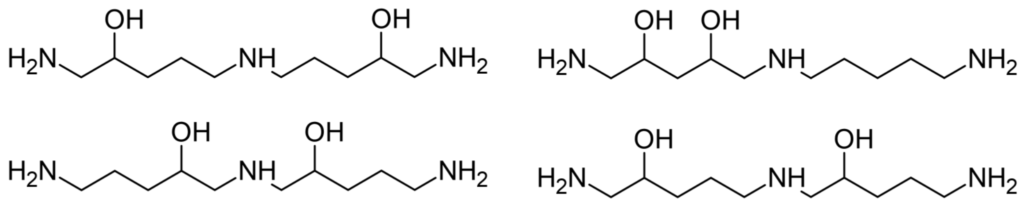

B

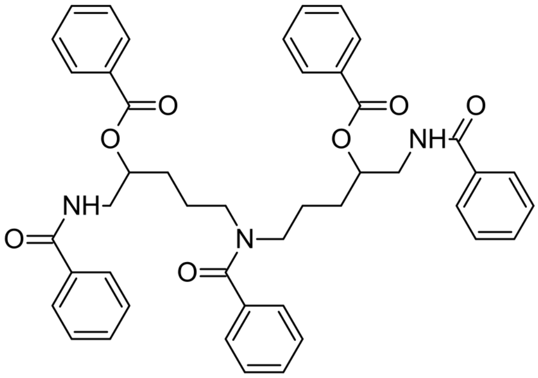

Molecular Formula: $C_{45}H_{45}N_3O_7$

Monoisotopic mass: 739.325751 Da

C

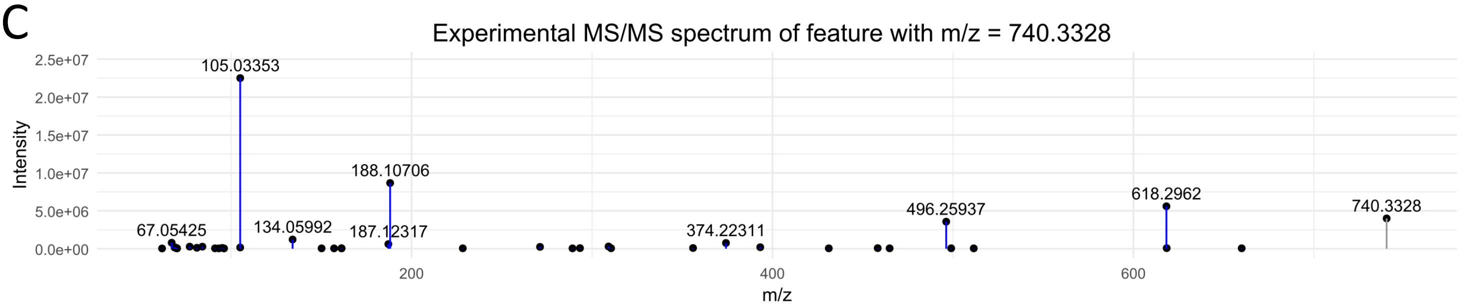


D

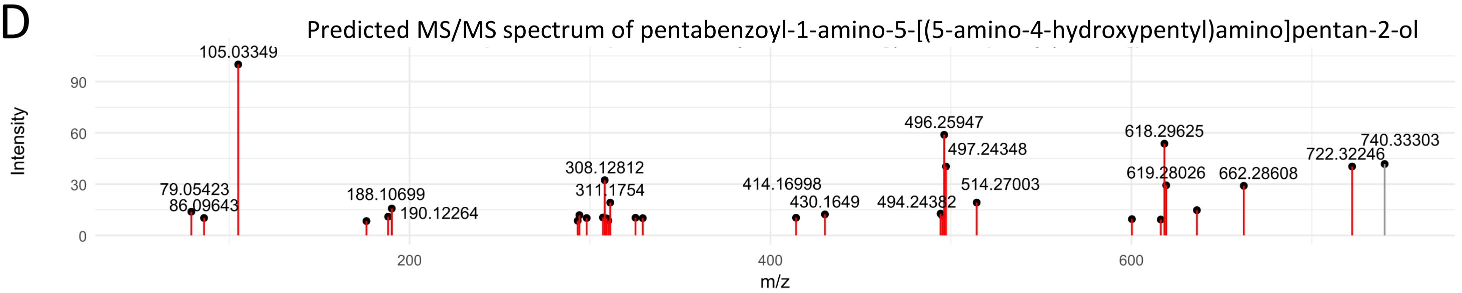


**Figure S5. MS/MS spectra of benzoylated features detected in cell extracts of *P. caledonica***. A. Possible structures of compound 3 (1-amino-5-[(5-aminopentyl)amino]pentane-diol) consistent with observed MS/MS spectra. B. Structure and molecular properties of benzoyl derivatives of compound 3, based on one of the likely structures shown in panel A. C. MS/MS profile of a feature with m/z= 740.3328 detected in derivatized extracts of cultures of *Paraburkholderia caledonica* SM2 pSEVA::*pavAB* grown at 16°C in LB medium. C. Predicted MS/MS spectrum of pentabenzoyl-1-amino-5-[(5-amino-4-hydroxypentyl)amino]pentan-2-ol determined using the CFM-ID web server with the SMILES notation: O=C(c1ccccc1)NCC(OC(=O)c2ccccc2)CCCN(C(=O)c3ccccc3)CCCC(OC(=O)c4ccccc4)CNC(=O)c5ccccc5. Peaks corresponding to precursor ions are shown in grey. Experimental MS data were analyzed in MZmine 4.6.1. Spectra data were exported and processed with the ggplot2 R package to generate plots. Note that the precise positions of the 2 hydroxy groups are uncertain.

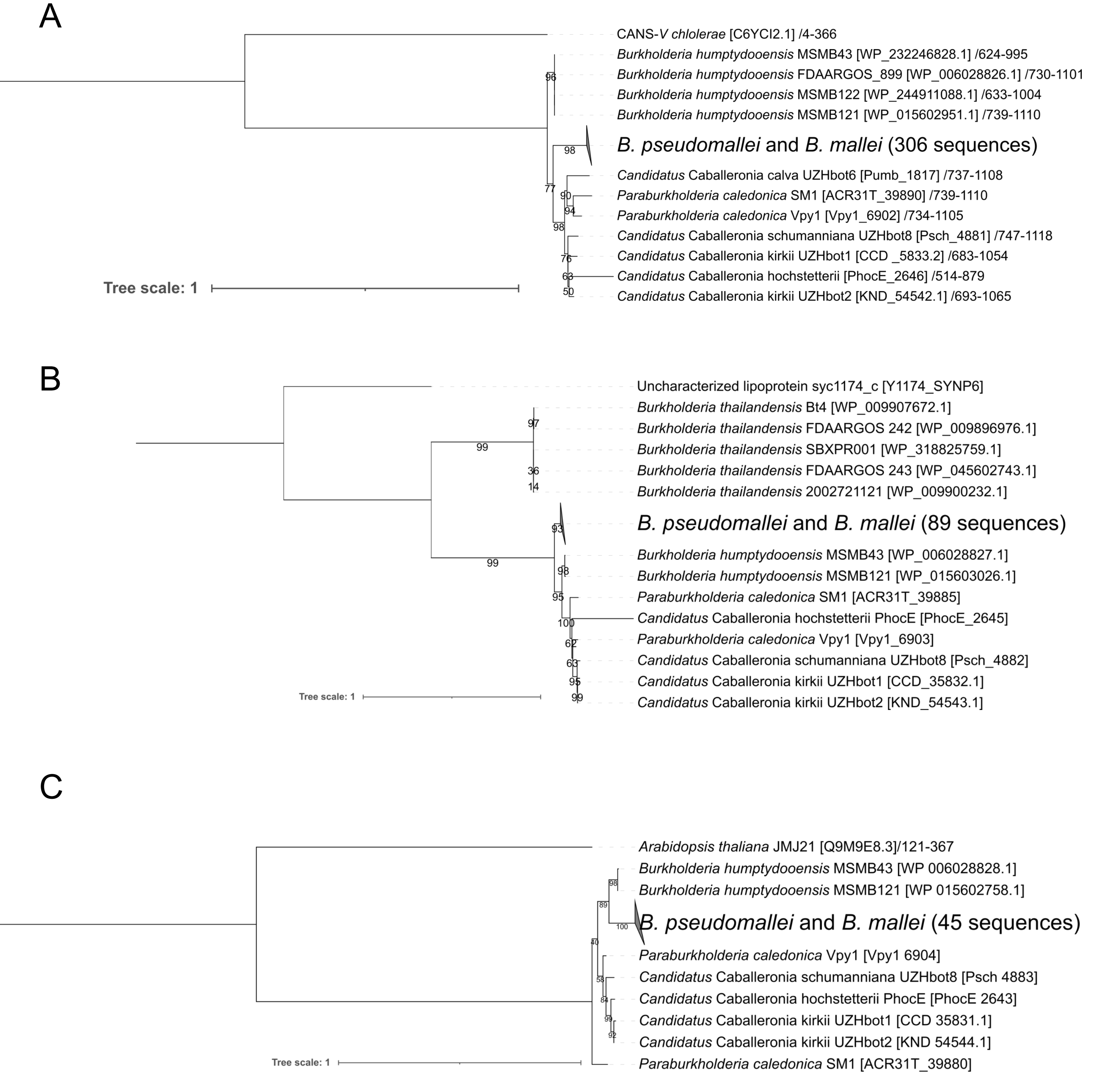


**Figure S6. Phylogenetic analysis of homologs of PavABC in the Burkholderia pseudomallei group.** A. Phylogenetic analysis of the putative decarboxylase C-terminal domain of PavA homologs. Numbers after the \ symbol indicate aligned positions relative to the initial protein sequence. B. Phylogenetic analysis of PavB homologs. C. Phylogenetic analysis of PavC homologs. Complete genomes and proteomes were downloaded with the NCBI datasets tool, using the taxon ID 111527 (Burkholderia *pseudomallei* group, 2096 genome sequence files). Proteomes were searched using BlastP and the PavA, PavB and PavC sequences of *P. caledonica* SM1. Hits were retrieved with an ad-hoc Python script, and identical proteins were removed prior to alignment with the SeqKit rmdup tool. Sequences were aligned with MAFFT and trimmed with Trimal in 'automated1' mode. To remove potential ambiguities in the alignment, the alignment of PavA was further trimmed manually to consider only the C-terminal decarboxylase domain (positions 675-1012 of the *P. caledonica* SM1 protein). Phylogenetic trees were generated with IQtree2 using the best fitting of the JTT, WAG or LG model families. Branch support values are from 1000 ultrafast bootstrap replicates. Protein accession numbers are indicated between brackets when available. To improve readibility, branches with average lengths below 0.01 are collapsed. Best blast hits in the SwissProt database were used to root the trees.

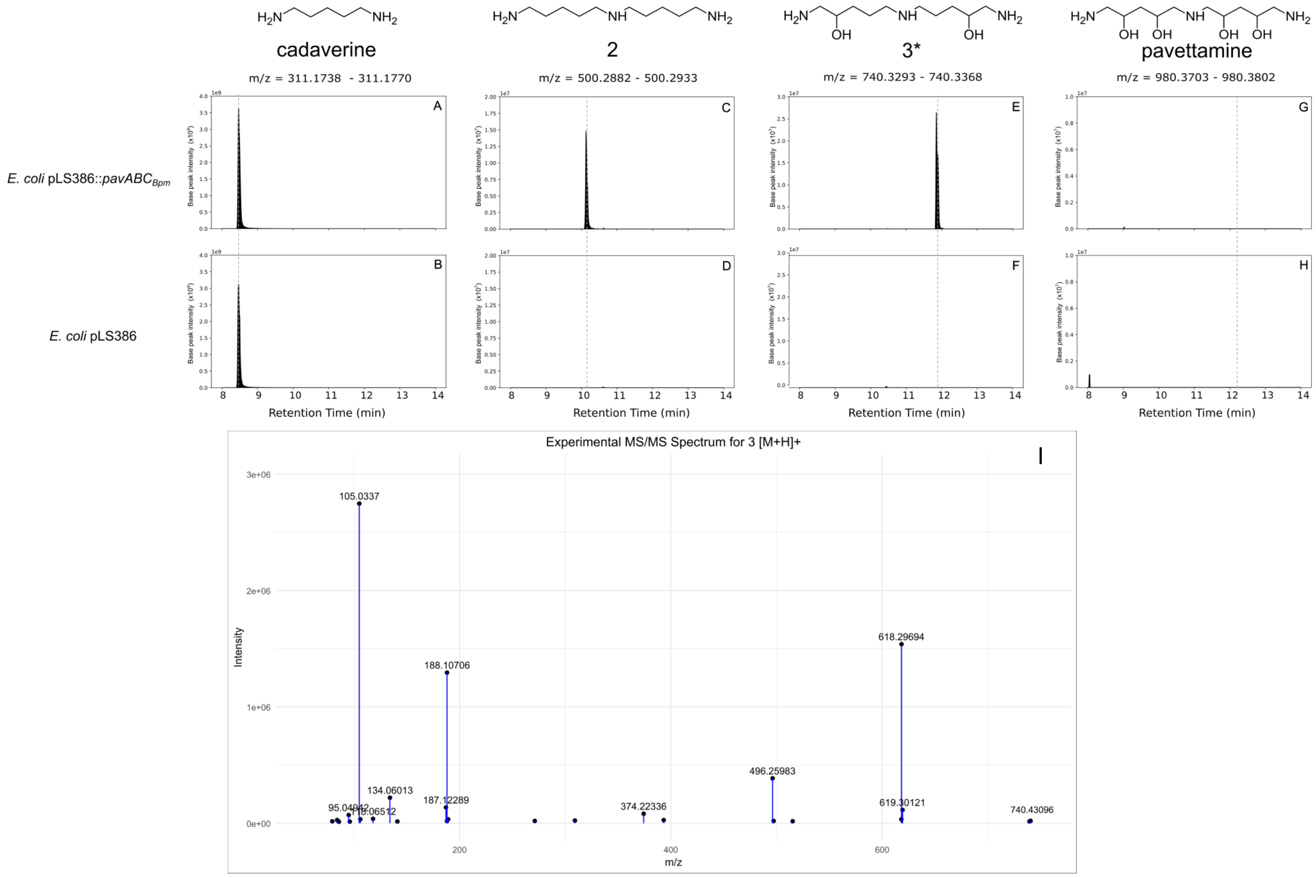


**Figure S7. Detection of pavettamine and putative pathway intermediates in cultures of *E. coli* expressing a codon-optimized *B. pseudomallei pavABC* operon.** A, B: Extracted ion chromatograms corresponding to the expected m/z range of the $[M+H]^+$ adducts of benzoyl derivatives of cadaverine. A, chromatogram of culture extracts of *E. coli* pLS386::*pavABC*$_{Bpm}$, and B, *E. coli* pLS386. C,D: Extracted ion chromatograms corresponding to the expected m/z range of $[M+H]^+$ adducts of benzoyl derivatives of compound 2. C, chromatogram of culture extracts of *E. coli* pLS386::*pavABC*$_{Bpm}$, and D, *E. coli* pLS386. E, F: Extracted ion chromatograms corresponding to the expected m/z range of the $[M+H]^+$ adducts of benzoyl derivatives of compound 3. E, chromatogram of culture extracts of *E. coli* pLS386::*pavABC*$_{Bpm}$; F, *E. coli* pLS386; G, H: Extracted ion chromatograms corresponding to the expected m/z range of the $[M+H]^+$ adducts of benzoyl derivatives of pavettamine. G, , chromatogram of culture extracts of E. *coli* pLS386::*pavABC*$_{Bpm}$; H, *E. coli* pLS386. Values indicated above the plots correspond to a 5 ppm interval around the theoretical m/z of the compounds. Grey dashed lines indicate the expected retention time of each analyte. I. MS/MS spectrum of a feature with m/z = 740.3331 and retention time = 11.74 min detected in cell extracts of E. *coli* pLS386::*pavABC*$_{Bpm}$, corresponding to the expected m/z and retention time of compound 3.
