## Supplementary material for "A fatal plant toxicosis described more than a century ago is caused by a bacterial endophyte": Table S1

**Table S1. Strains and plasmids**

| Strain | Description | Source |
| --- | --- | --- |
| FH1 | *Paraburkholderia caledonica*, wild-type*.* Isolate from leaves of *Fadogia homblei* collected in South Africa. | This study |
| SM1 | *P. caledonica* R-82532. Spontaneous nalidixic acid resistant mutant. Differs from strain R-82532 by a single T -> C mutation in the *gyrA* gene at position 3418872 of chromosome 2. Nal^R^ | This study |
| SM2 | *P. caledonica* SM1 *ΔpavABC*, Nal^R^ | This study |
| Top10 | *Escherichia coli*. K12 *ΔlacX74 araΔ139 Δ(ara-leu)*. General cloning host. | ThermoFisher Scientific |
| TG1 | *E. coli* K12 Δ(lac-pro),supE,thi,hsdD5, (F-traD36, proA^+^B^+^,lacI^q^,lacZΔM15). General cloning host. | (Carter et al. 1985) |
| DB3.1 *λpir* | E. coli *F^-^ gyrA462 endA1 glnV44 Δ(sr1-recA) mcrB mrr hsdS20 (r_B_^-^ m_B_^-^) ara14 galK2 lacY1 proA2 rpsL20 xyl5 Δleu mtl1 pir+*.  Host strain for pSMW2 and derivatives. | Thermo Fisher Scientific |
| ER2566 | *E. coli* host for protein expression. | (Fomenkov et al. 2017) |
| Plasmid |  |  |
| pRK2073 | Helper plasmid for tri-parental conjugation | (Leong et al. 1982) |
| pSNW2 | Suicide vector with I-SceI dependent counterselection. *oriR6K, Km^R^* | (Volke et al. 2021) |
| pQURE6 | Plasmid for conditional expression of I-SceI endonuclease. *Gm^R^* | (Volke et al. 2021) |
| pSEVA2313 | Broad host range expression vector; *OriV*(pBBR1), *P_EM7_*, Km^R^ | (Martínez-García et al. 2020) |
| pSEVA2313::*pavABC*  or pSEVA2313::ACR31T_39880-90 | *P. caledonica* SM1 ACR31T_39880-90 operon cloned downstream of P_EM7_ into a pSEVA2313 backbone | This study |
| pSEVA2313::*pavAB* | Plasmid derived of pSEVA2313::*pavABC* by targeted deletion of the *pavC* gene. | This study |
| pSEVA2313::*pavBC* | Plasmid derived of pSEVA2313::*pavABC* by targeted deletion of the *pavA* gene. | This study |
| pSEVA2313::*pavAC* | Plasmid derived of pSEVA2313::*pavABC* by targeted deletion of the *pavB* gene. | This study |
| pSEVA2313::*pav_bpm_*ABC | Synthetic gene construct reproducing the AA sequence of *B. pseudomallei* K96243 *pav_Bpm_ABC* cloned downstream of PEM7 | This study |
| pLS386 | Expression vector derived from pSMG259. | Laurent Sauviac (LIPME) and (Hoff et al. 2016) |
| pLS386::*pav_Bpm_ABC* | Synthetic gene construct reproducing the AA sequence of *B. pseudomallei* K96243 *pav_Bpm_ABC* cloned into the pLS386 expression vector. | This study |
