## Supplementary material for "A fatal plant toxicosis described more than a century ago is caused by a bacterial endophyte": Table S2

**Table S2. List of oligonucleotides used in this study.**

| **Primer** | **Sequence** | **Used for** |
| --- | --- | --- |
| Pcal_pavettamine_R-XbaI | GGGTCTAGACGCCGCTTTTTTTTGCGA | PCR amplification of *P. caledonica* FH1 *pavABC* operon. Construction of plasmid pSEVA2313::*pavABC* |
| Pcal_pavettamine_F-BamHI | GGGGGATCCAGCCGACCGAGAGACAA |  |
| pavABC_LeftP_BsaI_F | ACAGGTCTCGTACCCGGAGTTGATCTGGTGACC | PCR amplification of flanking sequences and construction of *P. caledonica* SM2 strain |
| pavABC_LeftP_BsaI_R | ACAGGTCTCGCATTCACGTGTTCCCTTGC |  |
| pavABC_RightP_BsaI_F | ACAGGTCTCGAATGTGACCCGCATATTCCCGAAAC |  |
| pavABC_RightP_BsaI_R | ACAGGTCTCGCAGGCCGGATCAGCGTCAACAACTG |  |
| 537_deletion_pavA_F | GGGGCTAGCATGCATGTCTATGAAAAGGC | Construction of pSEVA2313::pavBC |
| 538_deletion_pavA_R | GGGGCTAGCCCATTCACGTGTTCCCTTGC |  |
| 539_deletion_pavB_F | GGGGCTAGCTAACATGAAGAGAGTCGAAC | Construction of pSEVA2313::pavAC |
| 540_deletion_pavB_R | GGGGCTAGCTCATAGACATGCATGATTGAGC |  |
| 541_deletion_pavC_F | GGGGCTAGCTGACCCGCATATTCCCGAAAC | Construction of pSEVA2313::pavAB |
| 542_deletion_pavC_R | GGGGCTAGCCATGTTACGCCACCTGCACG |  |
| pLS386_Fw | CAGAAGACGAGAATTCCATGGATCCGCTAGC | PCR amplification of plasmid pLS386; construction of plasmid pLS386::*pavABC_Bpm_* |
| pLS386_Rev | CAGAAGACGATACCTCCTAAAAGTTAAACAAAAT |  |
| Bpse_pav_Fw | CAGAAGACGAGGTAGCGTAAATGGATCATCCTGATG | Construction of pSEVA2313::*pav*ABC*_bpm_*  and pLS386::*pav*ABC*_bpm_* |
| Bpse_pav_Rev | CAGAAGACGAATTCTTATTTAACCCACAGCGGATG | PCR amplification of *pavABC* from pSEVA2313::*pav*ABC*_bpm_*  And pLS386::*pav*ABC*_bpm_* |
